# SAR1 paralogs differ biochemically in assembly of the COPII coat

**DOI:** 10.1101/2020.02.09.940833

**Authors:** David B. Melville, Sean Studer, Randy Schekman

**Affiliations:** Department of Molecular and Cell Biology, Howard Hughes Medical Institute, University of California, Berkeley, Berkeley CA 94720

**Author notes:** Centroid Vaccines Inc., 3030 Bunker Hill St, Ste 115b, San Diego, CA 92109-5754. To whole correspondence should be addressed: Randy Schekman: Department of Molecular and Cell Biology, University of California at Berkeley, 475D Li Ka Shing Center #3370, Berkeley, CA 94720-3370.

**Keywords:** SAR1A, SAR1B, SEC23, SEC31, COPII, GTPase, apolipoprotein, intracellular trafficking, membrane trafficking, secretion

## Abstract

COPII-coated vesicles are the primary mediators of vesicular traffic from the ER to the Golgi apparatus. SAR1 is a small GTPase, which, upon GTP binding, recruits the other COPII proteins to the ER membrane. In mammals, there are two SAR1 paralogs which genetic data suggest may have distinct physiological roles, e.g. in lipoprotein secretion for SAR1B. We identified two clusters of amino acids that have conserved, paralog-specific sequences. One cluster is adjacent to the SAR1 GTP-binding pocket and alters the kinetics of GTP exchange. The other cluster is adjacent to the binding site of COPII components SEC31 and SEC23. We found that the latter cluster confers a SEC23A binding preference to SAR1B over SAR1A. In contrast to SAR1B, SAR1A is prone to oligomerize on a membrane surface. Importantly, in relation to its physiological function, SAR1B, but not SAR1A, can compensate for loss of SAR1B in lipoprotein secretion. The SEC31/SEC23-binding site-adjacent divergent cluster is critical for this function. These data identify the novel paralog-specific function for SAR1B, and provide insights into the mechanisms of large cargo secretion and COPII related diseases.

## Introduction

ER-to-Golgi protein trafficking is a key checkpoint in the sorting of proteins for secretion. Approximately one-third of all proteins are first assembled in the ER and then sorted to other destinations by COPII. Of the five core COPII proteins, the first to arrive at the ER membrane is SAR1. SAR1 is a small GTPase with an amphipathic helix that inserts into the membrane when SAR1 is in the GTP-bound state. SAR1 then recruits the remainder of the COPII complex, SEC23/24 and SEC13/31 heterodimers, to the ER membrane (1–5). It is thought that regulation of the GTPase activity of SAR1 is important for large cargo selection (6). Thus, SAR1 has two important roles: Recruitment of the other COPII proteins, and controlling the timing of COPII budding with its GTPase cycle.

Given its essential nature, it is perhaps unsurprising that SAR1 is extremely well conserved throughout evolution. The protein sequence of yeast and human SAR1 is either identical or strongly similar for ∼80% of the amino acid sequence, despite the fact that humans have greatly different secretory requirements from yeast. One area of divergence is that many invertebrates have only one paralog of SAR1, whereas mammals, and most vertebrates, have two. It is possible that these two paralogs have evolved unique functions to compensate for the diverse secretory needs of different cell types.

Genetic data provides some evidence that the two SAR1 paralogs, SAR1A and SAR1B, have divergent roles. For example, Loss of SAR1B leads to Chylomicron retention disease/Anderson’s disease (CMRD), which results in an inability to transport newly-synthesized chylomicrons out of intestinal epithelial cells (7–12). A similar phenotype was observed in zebrafish with loss of SAR1B (13), but not the more SAR1A-like SAR1AB. In cell culture, SAR1B knockout in the chylomicron secreting Caco-2/15 cells disrupts lipid homeostasis and induces oxidative stress, and inflammation(14, 15). SAR1A disruption produces a similar phenotype, but to a lower degree than SAR1B (14). Taken together, these data suggest the SAR1 paralogs have both overlapping and unique functions in cells, and likely differ biochemically, however little is known about the biochemical differences between SAR1A and SAR1B.

Here we identify two divergent clusters of conserved sequence differences between the two SAR1 paralogs. We find that a GTP-adjacent cluster alters GTP loading activity and direct interactions with SEC31A. We find that a second cluster in an apical α-helix causes SAR1B to bind more efficiently to SEC23 and SAR1A to homodimerize. We find that the apical α-helix is necessary and sufficient for the more efficient rescue of lipoprotein secretion by SAR1B than SAR1A. These data present clear biochemical differences between the two paralogs that provide a possible explanation for the differences seen in genetic data.

## Results

### SAR1A and SAR1B have two clusters of divergent amino acids

There are only 20/198 divergent amino acids that distinguish human SAR1A and SAR1B. Because highly conserved amino acid residues tend to be functionally important (16) we first wanted to compare how the SAR1A/B divergence appears in evolutionary history, and determine which divergent amino acids are most conserved. We retrieved the amino acid sequences from the ensembl database (17) and compared the sequences using clustal omega (18). We found that in reptiles, birds, and mammals there were conserved distinct SAR1A and SAR1B paralogs, whereas in some fish, such as zebrafish, there was a distinct SAR1B allele and a more intermediate SAR1AB allele closer to the invertebrate ancestral gene (Figure 1A). These data suggest that comparing mammalian, reptile, and bird alleles would provide a broad consistent background for determining which paralog-specific amino acid differences are conserved.

**Figure 1:**
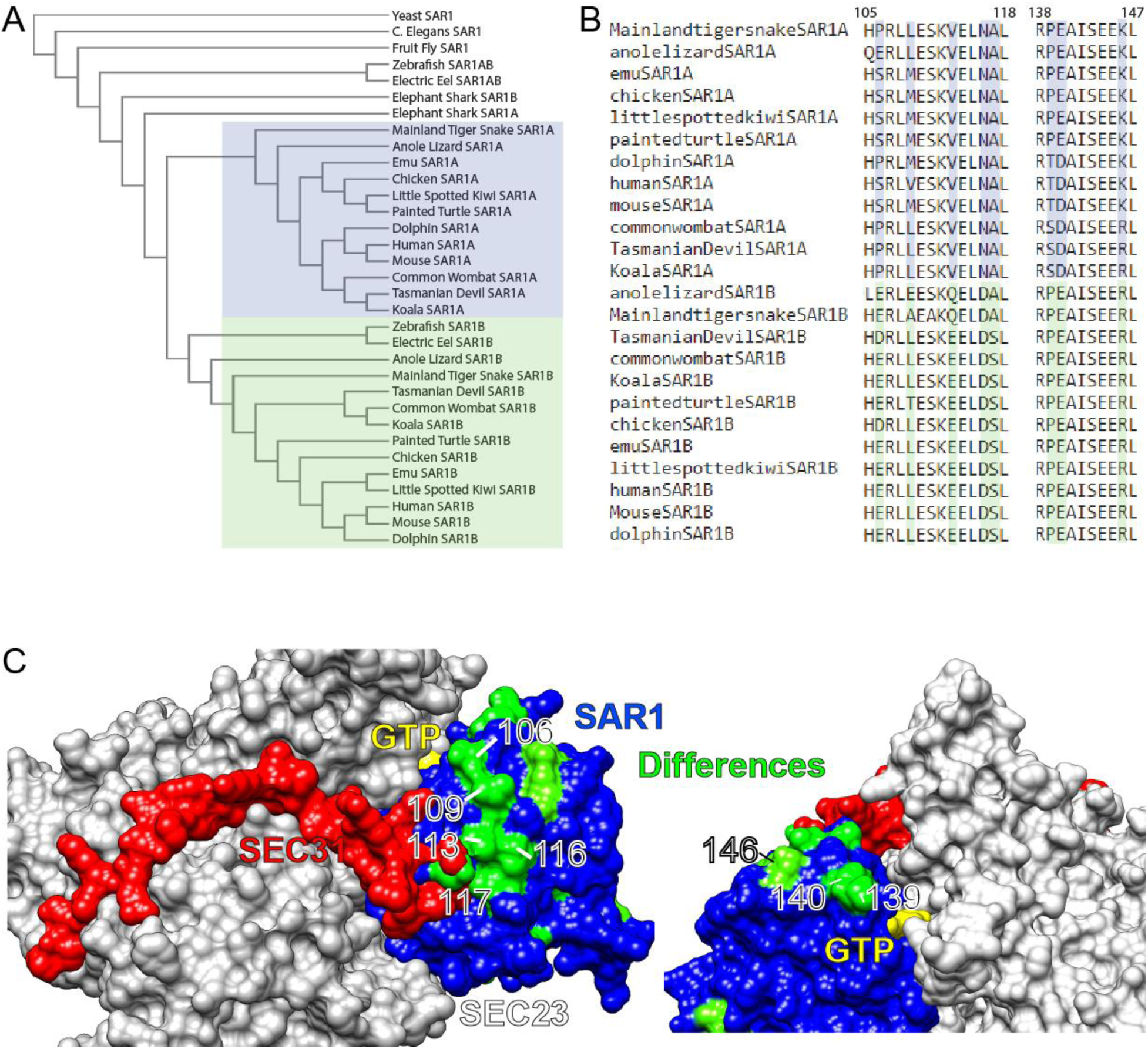
The evolutionary conservation of SAR1 paralogs. (A) Phylogenetic guide tree of SAR1 paralogs in three species each of invertebrates, fish, reptiles, birds, marsupials, and non-marsupial mammals. (B) Sequence alignment of apical α-helix and GTPase-adjacent clusters of divergent amino acids. (C) Structure of human SAR1A and B modeled onto yeast SEC23/SAR1/SEC31 with divergent amino acids highlighted in green.

We next looked for candidate amino acids that might lead to divergent functions. Primary sequence alignment of human SAR1A and SAR1B found three clusters where three or more amino acids diverged in close proximity. Of those three clusters, two were highly conserved among mammals, reptiles, and birds (106-117 and 139-146) (Figure 1B), whereas one (162-164) was less conserved.

SAR1 residues 139-146 are adjacent to the GTP-binding pocket and residues 106-117 are on an α-helix near the known binding site of SEC31 on SEC23 (Figure 1C). Notably, the divergent amino acids on the α-helix all appear on the exposed surface of the protein, suggesting that they may have a role in protein-protein interactions. Notably, the importance of the interaction between SEC31 and the SAR1 GTPase cycle for large cargo secretion has been well documented (6, 19–21). We hypothesized that the GTP-adjacent cluster of divergent residues may play a role in either GTP exchange or hydrolysis, whereas the SEC23/31-adjacent divergent helix may play a role in binding SEC31.

### SAR1A has faster GTPase exchange than SAR1B

In order to test whether the divergent GTP-adjacent amino acids led to different GTP cycle activity in SAR1, we utilized a tryptophan fluorescence-based assay performed with purified human proteins (22–24). In this assay, the nucleotide-bound state of Sar1 is monitored by relative fluorescence measurements. The intrinsic tryptophan fluorescence of SAR1-GTP is significantly higher than that of SAR1-GDP.

We first analyzed the kinetics of GTP exchange upon addition of GTP to the reaction. We found that with full-length SAR1 in the presence of liposomes, SAR1A loaded significantly faster than SAR1B (Figure 2A). This difference applied to soluble forms of SAR1 that lack the N-terminal amphipathic helix and as a result do not require liposomes (Figure 2B). Addition of the SAR1 guanine exchange factor (GEF) SEC12 increased the loading speed of both paralogs proportionally (Figure 2C). We confirmed that this effect was unrelated to hydrolysis by substitution of a non-hydrolyzable GTP analog GMP-PNP (Figure 2D).

**Figure 2:**
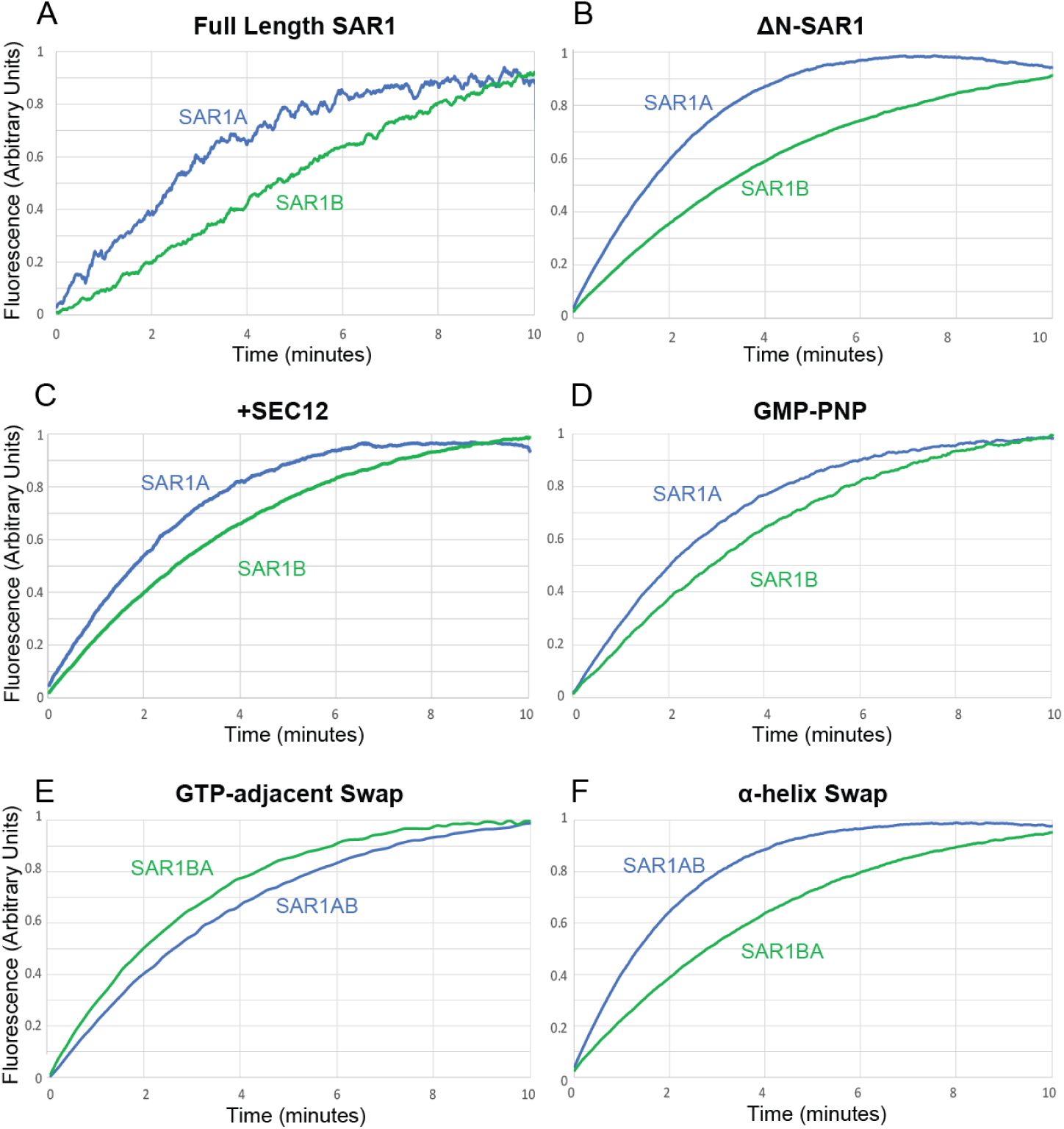
GTP exchange in SAR1A and SAR1B. (A-F) Tryptophan fluorescence assay measuring loading of GTP into SAR1A and SAR1B with (A) Full-length SAR1 and liposomes, (B) ΔN-SAR1, (C) ΔN-SAR1 and SEC12, (D) ΔN-SAR1 and non-hydrolyzable GMP-PNP in place of GTP, (E) ΔN-SAR1 with GTP-adjacent divergent amino-acid cluster swapped, (F) ΔN-SAR1 with SEC31-binding site adjacent divergent amino acid cluster swapped.

We hypothesized that this difference may be due to the GTP-adjacent divergent residues. To test this, we purified SAR1 proteins with the divergent amino acids swapped between paralogs (SAR1A>B-GTPa and SAR1B>A-GTPa). We found that changing these three amino-acids reversed the kinetic differences between SAR1A and SAR1B (Figure 2E). Conversely, we found that constructs in which the amino acids of the divergent apical α-helix (SAR1A>B-helix and SAR1B>A-helix) were swapped did not reverse the kinetics (Figure 2F). These data suggest that the GTP-adjacent divergent residues are necessary and sufficient for the increased kinetics of SAR1 GTP exchange.

Having observed differences in GTP loading, we probed the SAR1 paralogs for GTP hydrolysis using the tryptophan-fluorescence assay. We did not find a strong consistent difference between the two paralogs in the presence of either different sizes of liposomes (Supplemental Figure 1A) or SEC23A or SEC23B (Supplemental Figure 1B). This evidence suggests that although SAR1A exchanges nucleotide more quickly that SAR1B, GTP hydrolysis may be unaffected, at least *in vitro*.

### SAR1A binds the GTPase activating fragment of SEC31 more strongly that SAR1B

One of the two divergent clusters between SAR1A and SAR1B is adjacent to the known SEC31/SEC23 binding site. We therefore hypothesized that this cluster may be important for direct binding of SEC31 to SAR1. To test the extent of direct binding between SAR1 and SEC31, we utilized a liposome flotation assay. Purified recombinant human SAR1 and the GTPase activating fragment of SEC31A (SEC31A-af) were incubated with synthetic liposomes and GMP-PNP. The reaction was applied to the bottom of a sucrose density gradient. After a high-speed centrifugation step, liposomes carried bound SAR1, and any SEC31 bound to that SAR1, with them as they floated to the top of the sucrose gradient (Figure 2A). We then evaluated the levels of SAR1 and SEC31A-af present on the liposomes by SDS-PAGE followed by SYPRO-Ruby staining.

We found that SAR1A was approximately 2-fold more efficiently recruited to SEC31A-af than SAR1B (Figure 2B,C). When we performed the same assay with SAR1A>B-helix and SAR1B>A-helix, however, the difference became negligible, suggesting that the divergent α-helix played a role in SEC31 binding, but was not the sole reason for differences between SAR1A and SAR1B.

Here we made a secondary observation. Despite having similar molecular weights, SAR1B migrated more slowly in SDS-PAGE gels than SAR1. However, SAR1A>B-helix migrated similarly to SAR1B, and SAR1B>A-helix similarly to SAR1A (Figure 2B). Therefore, whatever causes this different migration of the two paralogs is contained in the α-helix.

We then compared the SAR1A>B-GTPa and SAR1BA-GTPa. Unexpectedly, swapping the GTPase-adjacent divergent residues had a more significant effect than swapping the α-helix (Figure 3 B,C), suggesting that the GTP-adjacent residues have as much a role, if not more, than the divergent α-helix in SEC31 binding.

**Figure 3:**
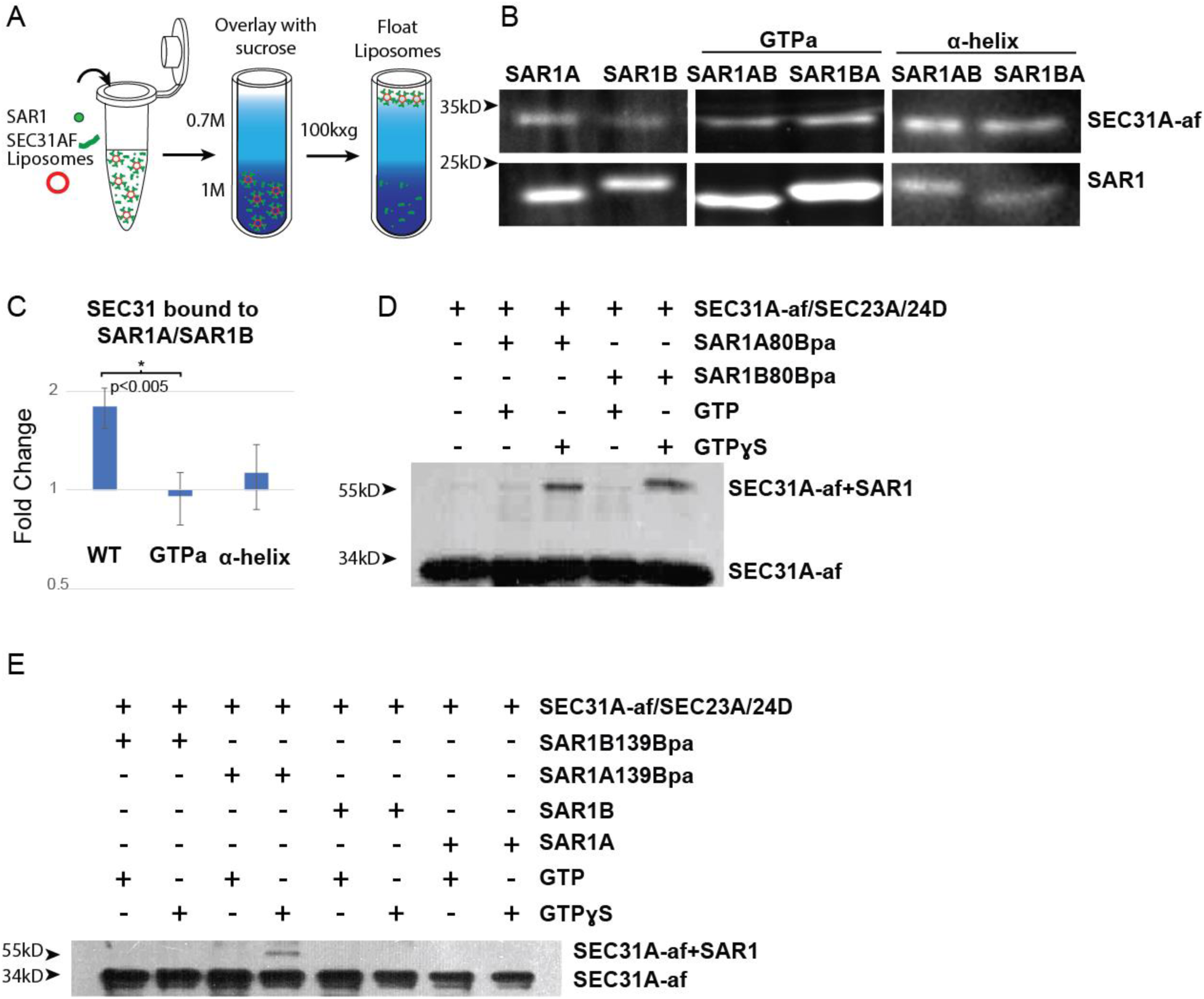
Binding of SAR1 paralogs and the SEC31A GTPase-activating fragment. (A) Schematic of liposome flotation assay, where COPII proteins are incubated with liposomes, then floated through a sucrose gradient to determine which proteins bind to liposomes. (B) SDS-PAGE followed by Sypro ruby stain of SEC31A-af recruited by SAR1 to floated liposomes. (C) Quantification of liposome flotation experiments (N>3 for each condition). (D,E) Photo-crosslinking assay of SAR1 with unnatural amino acid at position 80 (D) and 139 (E) incubated with SEC31A-af and photoactivated. Upper band indicates crosslinked SAR1/SEC31A-af.

To confirm whether the GTP-adjacent residues directly bind SEC31A-af, we used photo-crosslinking to an unnatural amino acid in recombinant SAR1 protein. We verified that the assay detected SEC31A-af binding by inserting an unnatural amino acid at position 80, directly adjacent to the known SEC23A/SEC31A-af binding site. Crosslinking was stimulated by incubation with the non-hydrolyzable GTP analog GTPγS and either SAR1 paralog (Figure 3D). We then repeated this with a GTP-adjacent residue, inserting the unnatural amino acid at position 139 (Threonine in SAR1A, Proline in SAR1B). Crosslinking was stimulated by GTPγS primarily in SAR1A (Figure 3D), consistent with our liposome flotation data. These data suggest that both divergent clusters play a role in SAR1-SEC31 binding, and that SAR1A has a higher affinity for direct binding of SEC31.

### SAR1B binds SEC23 more strongly than SAR1A

Although our data show that SAR1 can directly bind SEC31, SAR1 normally binds SEC31 in conjunction with SEC23. Using the liposome flotation assay, we compared recruitment of SEC23 and SEC31 by the two SAR1 paralogs. Because SEC23A and SEC23B also have paralog-specific roles in large cargo secretion, we also tested whether SAR1 had a different affinity for SEC23A or SEC23B by adding both in competition to the reaction.

We found a strong divergence between SAR1A and SAR1B in their recruitment of either paralog of SEC23. SAR1B recruits SEC23 at a ∼5 fold higher level than SAR1A (Figure 4A,C). Addition of SEC31A-af increased the binding of both SAR1 paralogs for both SEC23 paralogs, but the binding of SAR1B to SEC23 remained ∼5 fold higher than SAR1A.

**Figure 4:**
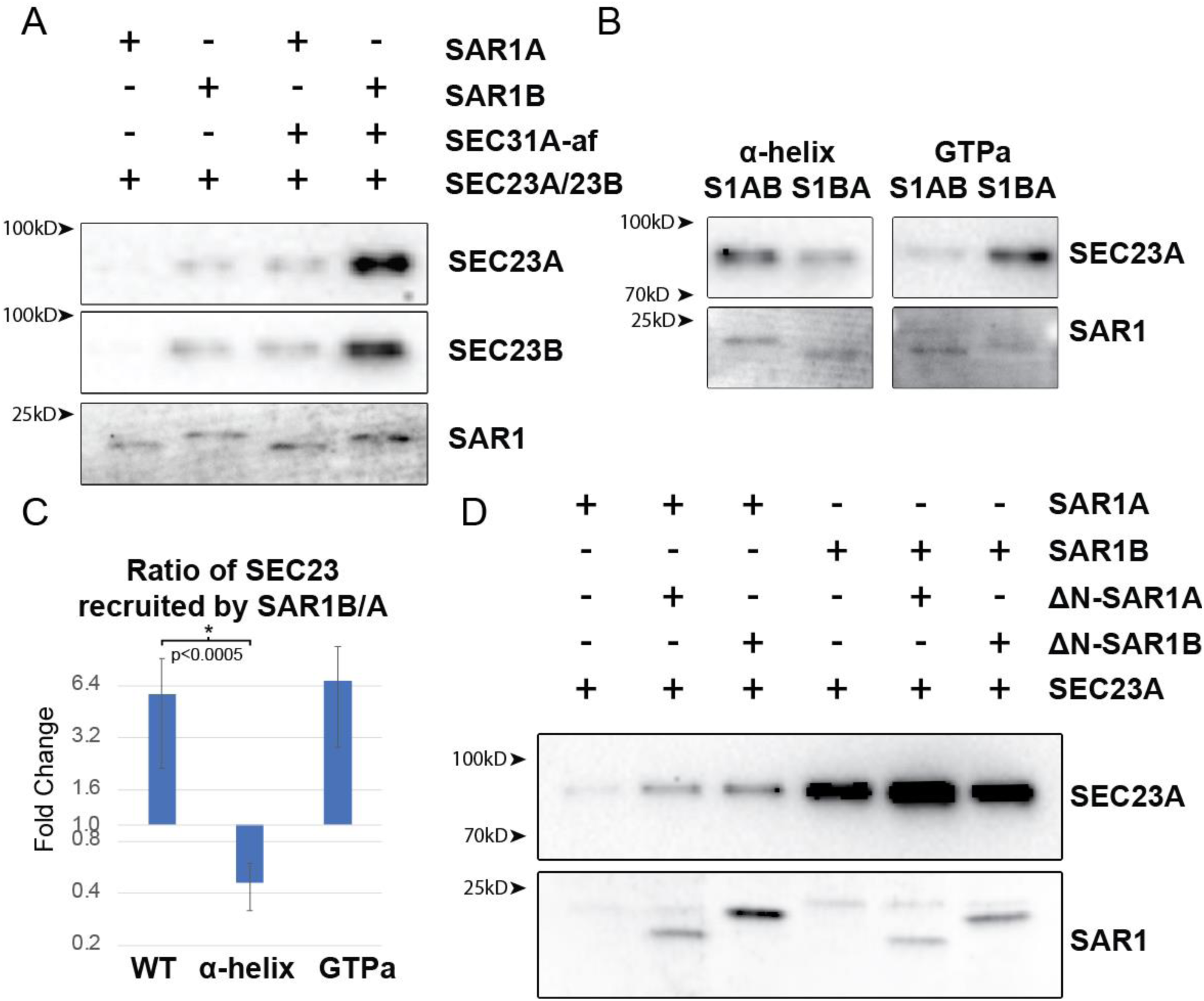
Binding of SAR1 and SEC23. (A-B) SDS-PAGE followed by Sypro ruby stain of SAR1 WT (A) or indicated divergent amino acid group swaps (B) and immunoblot of indicated SEC23 paralog recruited to liposomes by SAR1. (C) Quantification of liposome flotation experiments (N>3 for each condition). (D) SDS-PAGE followed by Sypro ruby stain of indicated SAR1 and ΔN-SAR1 recruited to liposomes by SAR1 and immunoblot of SEC23A recruited to liposomes by SAR1.

In order to determine which amino acids in SAR1 were responsible for the divergence in SEC23 recruitment, we repeated the flotation assay with the GTPa and helix SAR1 variants. We found that swapping the divergent α-helix reversed affinity for SEC23, reducing SAR1B>A-helix binding to SEC23 by ∼2 fold instead of a 5 fold increase (Figure 4B,C). These data suggest that the divergent helix leads to SAR1B having a higher affinity for SEC23.

We hypothesized that SAR1B’s greater affinity for SEC23 may interfere with SAR1A’s ability to recruit SEC23 and the remaining elements of the COPII coat if it were present in excess over SAR1A. To test this, we performed the liposome flotation assay using the soluble N-terminal deleted SAR1. We hypothesized that addition of soluble SAR1B may decrease the amount of SEC23 recruited by SAR1A by sequestering the available pool of protein. We found, however, the opposite: Addition of either soluble SAR1 paralog enhanced SEC23 recruitment (Figure 4D).

Enhanced recruitment of SEC23 may be due to the formation of SAR1 dimers on the membrane (25–27). In fact, we found a significant amount of soluble SAR1 recruited to the membrane, although we were unable to distinguish soluble SAR1B from SAR1A as the two have the same SDS-PAGE mobility. SAR1A appeared to be more prone to recruiting soluble SAR1 than SAR1B (Figure 4D). These data suggest that SAR1A may have a higher affinity for homodimerization than SAR1B.

### SAR1A homodimerizes more strongly than SAR1B

In order to evaluate the oligomerization of SAR1 on a membrane surface, we developed a liposome aggregation assay. We hypothesized that SAR1 dimerization could cause small liposomes to aggregate, decreasing the number of particles and increasing their size.

We prepared small 100nm synthetic liposomes and incubated them with SAR1 and GTP-PNP. After overnight incubation, SAR1A-containing liposomes formed aggregates easily visible by light microscopy (Figure 5A), whereas SAR1B-containing liposomes were indistinguishable from liposomes alone. The size and number of particles in the suspension were then evaluated with a Nanosight particle analyzer. In order to detect smaller particles that could be reliably quantified by Nanosight, we used lower protein concentrations and shorter incubation times. Under these conditions, SAR1A-containing liposomes were larger and fewer in number than the SAR1B containing liposomes (Figure 5B,C). We further found that, for both SAR1A and SAR1B containing liposomes, addition of SEC23 greatly increased the size of particles and reduced their number (Figure 5B), as would be expected because SEC23 binds directly to SAR1A and B.

**Figure 5:**
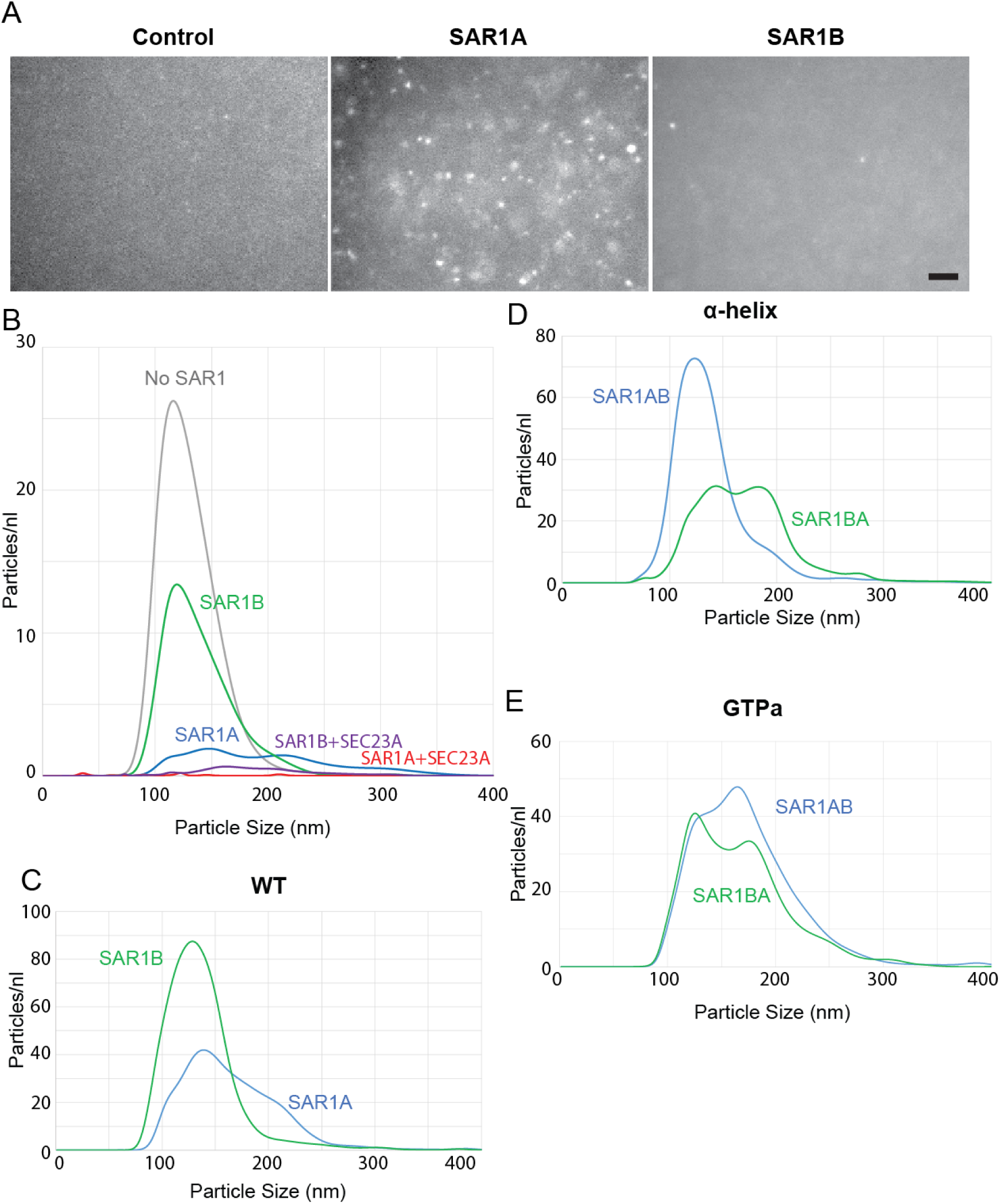
Liposome aggregation by SAR1 paralogs. (A) Fluorescent microscopy of 100nm liposomes incubated overnight at 37°C. Only aggregates >200nm can be clearly resolved. Scale bar 10μm. (B-E) Nanoparticle analysis of 100nm liposomes incubated with GMP-PNP and indicated proteins. Particle count inversely correlated to aggregation.

To test whether this effect was due to either of the divergent peptide clusters, we performed this assay with the GTPa and helix SAR1 constructs. We found that swapping the divergent helix caused SAR1BA-helix to promote liposome aggregation more than SAR1AB-helix (Figure 5D), suggesting that the divergent helix is the primary driver of SAR1A oligomerization/aggregation. We found that swapping the GPTa cluster also resulted in increased aggregation by SAR1BA-GTPa (Figure 5E). This effect is much milder than that seen with SAR1BA-helix, suggesting that the GTP-adjacent cluster may also play a role in oligomerization, but a relatively minor one compared to the apical helix. These data suggest that SAR1A oligomerizes on a membrane surface and the divergent helix has a more prominent role in this association than the GTP-adjacent cluster.

### The divergent helix in SAR1B facilitates rescue of lipoprotein secretion

In order to test the functionally significant differences of the SAR1 paralogs in cells, we measured apolipoprotein secretion in transfected cells. For this purpose, we developed CRISPR-Cas9-mediated SAR1B knockdown with the lipoprotein-secreting rat hepatoma cell line McArdle RH7777. Cells were incubated in oleic acid-containing medium from which samples were withdrawn every 1-3 h. Aliquots of medium were subjected to density sedimentation on an Optiprep gradient to collect the buoyant lipoproteins. Secreted APO1B was detected and quantified by immunoblot. As has been previously found (28), loss of SAR1B resulted in a substantial reduction of APOB100 in the medium (Figure 6A).

**Figure 6:**
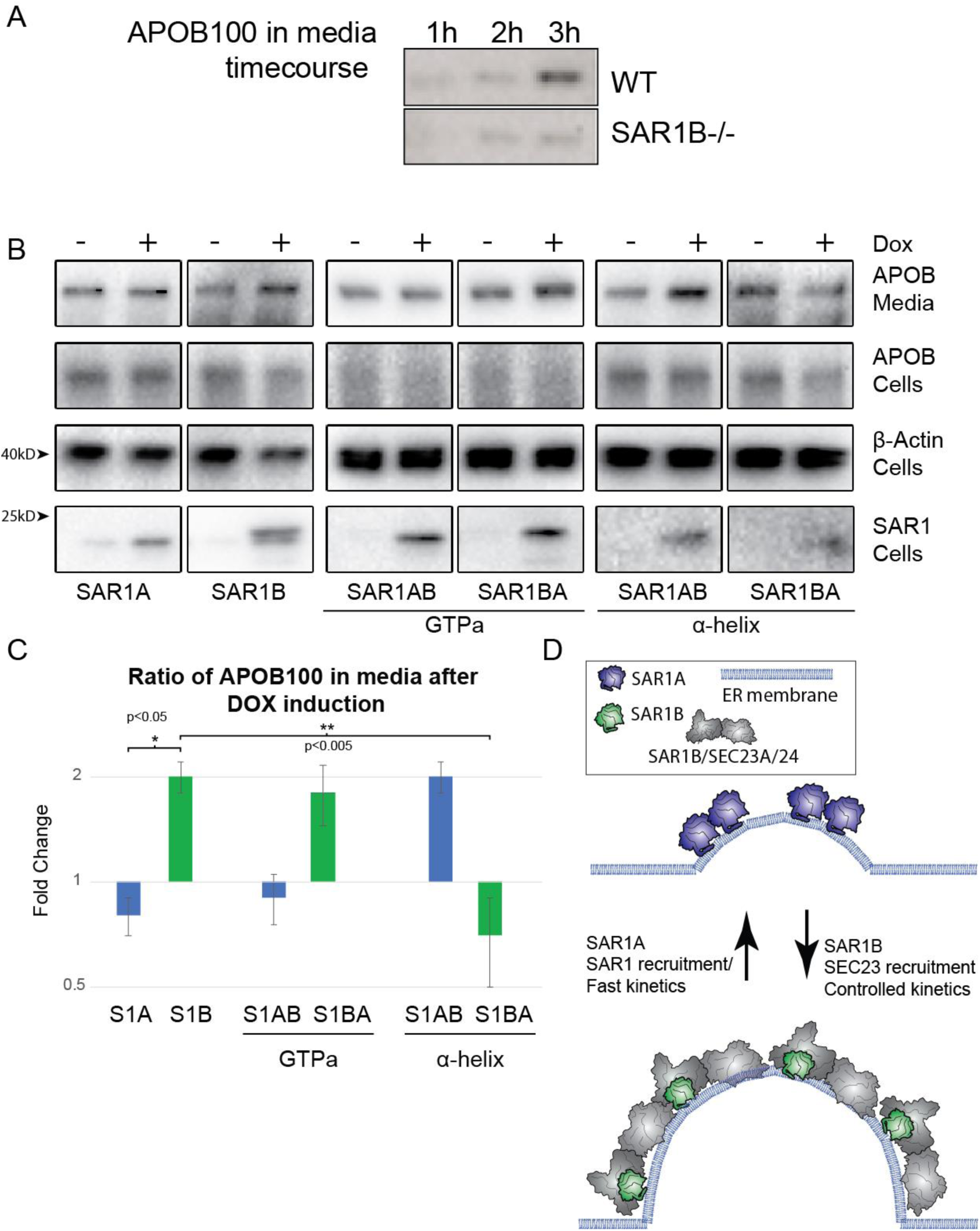
Apolipoprotein B 100 secretion and SAR1. (A) Immunoblot of APOB in top fraction of OptiPrep gradient from media of WT and SAR1B-/- McArdle cells incubated with oleic acid for the indicated time. (B) Immunoblot of APOB in SAR1B-/- McArdle cells transformed by lentivirus to overexpress indicated SAR1 constructs after doxycycline induction. β-Actin used as a loading control. (C) Quantification of APOB secreted into media (n=3). (D) Proposed model of divergent roles of SAR1 paralogs in cells. SAR1A proposed to homodimerize at the membrane for ERES remodeling, whereas SAR1B, which has a higher affinity for SEC23, leads to higher recruitment of SEC23/24 heterodimer, enabling better secretion of large cargos that rely on tighter control of the kinetics of COPII vesicle formation.

Using the lipoprotein secretion-deficient SAR1B-/- cells, we generated inducible cell lines using lentivirus and a tetracycline repressor system to control the expression of SAR1 by addition of doxycycline. We found that overexpression of SAR1B increased the amount of APOB100 secreted into the medium, whereas overexpression of SAR1A did not (Figure 6B). Overexpression of the GTP-adjacent swap SAR1B>A produced an approximately 2-fold increase in ApoB secretion, similar to the effect of wild-type SAR1B, suggesting that the GTP-adjacent amino acid cluster is not relevant to the SAR1B-specific function. Overexpression of the helix swap SAR1B>A failed to increase APOB100 secretion, whereas SAR1A>B increased ApoB secretion approximately 2-fold, suggesting that the helix is most important for the paralog-specific function of SAR1B in lipoprotein secretion.

Taken together, these data suggest that SAR1A and SAR1B differ biochemically. The paralogs have two divergent clusters of amino acids, one adjacent to the GTP binding pocket and one in an α-helix on the apical side of the protein, that have conserved-paralog specific sequences and functions. The GTPase-adjacent cluster causes SAR1A to exchange GTP more rapidly than SAR1B. The apical α-helix cluster causes SAR1B to have a higher affinity for SEC23A, and serves an important role in lipoprotein secretion.

## Discussion

Vertebrate cells have a wide variety of distinct secretory requirements, both in the types of cargo and the overall cargo load. Evolution of different paralogs of the basic machinery gives cells specialized tools to deal with the particular needs of a given cell at a given time. We have found how small changes in the primary sequence of SAR1 paralogs lead to different biochemical characteristics that have significant impacts on cellular function.

### SAR1 as a remodeler and cargo size

It is easy to think of SAR1 as just a small part of a bigger COPII machinery, rather than focusing on the important roles SAR1 itself plays. Hannah *et al.* (26) showed that the amphipathic helix of GTP-bound Sar1 stably penetrates the endoplasmic reticulum membrane, promoting local membrane deformation. As membrane bending increases, Sar1 membrane binding is elevated, ultimately culminating in GTP hydrolysis, which may destabilize the bilayer sufficiently to facilitate membrane fission. Lee *et al.* (29) showed that SAR1 promotes significant membrane remodeling by itself, a finding that has been amplified by compelling atomic-force microscopy videos of rapid rearrangement of membrane induced by SAR1. SAR1 dimerization may aid in membrane remodeling (25–27).

Recent cryo-tomography (30) of COPII assembled on membrane suggests quite the opposite situation. Their data present an elegant, ordered array of SAR1/SEC23/SEC24 subunits on a tubular membrane with no evidence of SAR1 dimerization in the assembled COPII structures. Hutchings *et al.* propose that COPII forms small vesicles using inner coat patches that insert Sar1 amphipathic helices randomly and curve the membrane in all directions, similar to what was seen with atomic-force microscopy. They propose that larger structures, by contrast, are formed by extensive assembly of the inner coat, consistent Sar1 orientation and parallel insertion of its amphipathic helix.

Taking these results together, and having identified biochemical differences between SAR1 paralogs, we propose the following model for how the SAR1 paralogs might play a role in cells (Figure 6D).

### A bimodal model of SAR1 and COPII behavior

Trafficking of large cargos presents a special challenge to the COPII machinery. Disruption of transcriptional regulators of COPII can also lead to large cargo-specific defects (31– 33), suggesting that large cargos are especially sensitive to changes to COPII dynamics. The kinetics of COPII assembly has been found to be especially important for trafficking of both collagen and large lipoproteins (20, 21, 34). In collagen trafficking, for example, TANGO1 competes with SEC31 for binding to SEC23, postponing the final steps of budding until collagen is loaded into vesicles (21, 34). Conversely, for small cargos, faster kinetics would allow cells to more quickly address their trafficking needs.

We propose that under conditions where speed is more critical than size, such as when small cargos needs to be trafficked, SAR1 dimers insert their amphipathic helix into to membrane, creating high levels of curvature, and thus recruit more SAR1 to the membrane both through dimerization and an affinity for highly curved membrane. This leads to numerous small vesicles and fast budding. This process may be most efficiently driven by SAR1A, with its fast GTP loading and efficient oligomerization (Figure 6C, top).

Under conditions where speed is less critical than size, such as large cargo secretion, SAR1 recruits SEC23/24 heterodimers and forms ordered lattices that allow for slower, more controlled, membrane curvature, and packaging of large cargos. This process may be most efficiently driven by SAR1B, with its high affinity for SEC23 and less efficient oligomerization (Figure 6C, bottom).

In this model, controlling the balance of the two SAR1 paralogs would give cells a mechanism to respond to different secretory requirements.

### Paralog-specific functions not unique to SAR1

The existence of multiple paralogs of mammalian COPII subunits provides an opportunity for fine tuning of cargo sorting dependent on the physiologic needs of different cells and tissues. In addition, COPII protein levels are dynamically regulated (31, 32). Loss-of-function of these paralogs leads to many distinct phenotypes (35–42). In an extreme example, SEC13 has a dual role as nuclear pore component as well as a subunit of the outer shell of the COPII coat (43).

The unique roles of COPII paralogs present a dynamic picture of COPII regulation of and response to protein trafficking and other cellular needs. Rather than a single complex with a single purpose, COPII paralogs provide a cellular membrane trafficking toolkit.

## Experimental Procedures

### Phylogenetic analysis

Protein sequences were aligned using ClustalOmega (https://www.ebi.ac.uk/Tools/msa/clustalo/). A cladogram tree was constructed using Clustal alignment.

### Antibodies

Commercially available antibodies used for immunoblotting were as follows: Goat anti-apolipoprotein B (EMD, Hayward, CA; #178467 1:500 for immunoblot) (Figure 6A) Rabbit anti-apolipoprotein B (Proteintech Rosemont, IL #20578-1-AP)(Figure 6B).

### Lentivirus production and adipocyte transduction

Human SAR1 was subcloned into pLenti-puro (Addgene Cambridge, MA; Plasmid #39481). The plasmid containing SAR1 was transfected with Lipofectamine 2000 (Invitrogen) following the manufacturer’s recommendations into HEK293T cells at 50% confluence the day of transfection along with lentiviral packaging plasmids pVSVg (3.5 µg) and psPAX2 (6.5 µg; Addgene). Transfection was performed using one 10-cm dish. After a 24-h transfection, the medium was changed, and after an additional 24 h, the medium was removed and filtered through a 0.45-µm low-protein binding membrane (VWR International, Radnor, PA). McArdle-RH7777 or IMR-90 were then transduced with the virus with 8 µg/ml polybrene (Sigma-Aldrich). After 24 h, medium was replaced with fresh medium, and after additional 24 h, 2 µg/ml puromycin (Sigma-Aldrich) was added to select transduced cells.

### Protein purification

Human SAR1 and SEC31-GTP activating fragment proteins were expressed in *Escherichia coli* and purified as GST-fusions and then cleaved, as described for hamster Sar1 purifications (44). In brief, a bacterial lysate was first centrifuged at 43,000xg for 15 min, then the supernatant fraction was further centrifuged at 185,000xg for 1 h. The supernatant was incubated with prewashed glutathione agarose (1 ml slurry/l bacteria; Thermo Fisher Scientific) for 1 h at 4°C. Agarose was washed with wash buffer (50 mM Tris, pH 7.4, 150 mM NaCl, 0.1% Tween, 5 mM MgCl_2_, and 100 µM GDP), and protein was eluted by cleaving with 20 U/ml thrombin (Roche) in TCB (50 mM Tris, pH 8, 250 mM KoAc, 5 mM CaCl_2_, 5 mM MgCl_2_, and 100 µM GDP). Human SEC23/24 paralogs and variants were purified from lysates of baculovirus-infected insect cells, as described previously (44). In brief, insect cell lysates were centrifuged at 185,000xg for 1 h and 30% ammonium sulfate was added to the supernatant fraction at 4°C. The precipitant was collected by centrifugation at 30,000xg for 30 min and solubilized in no-salt buffer (20 mM Hepes, pH 8, 10% glycerol, 250 mM sorbitol, 0.1 mM EGTA, 5 mM β-mercaptoethanol, and 10 mM imidazole). The solubilized 30% ammonium sulfate precipitant was cleared at 30,000xg for 20 min, and the supernatant was incubated with prewashed Ni-NTA resin (1.25 ml slurry/l insect cells; Thermo Fisher Scientific) for 1 h at 4°C. Ni-NTA was washed with 20 mM Hepes, pH 8, 10% glycerol, 250 mm sorbitol, 500 mM KoAc, 0.1 mM EGTA, 5 mM β-mercaptoethanol, and 50 mM imidazole and eluted with 250 mM imidazole. Ni-NTA–eluted SEC13/31A protein was further purified using an anion exchange column (MonoQ) on an AKTA FPLC system (GE Healthcare).

### Immunoblotting

Standard immunoblotting procedures were followed. In brief, samples were resolved on 4–20% polyacrylamide gels (15-well, Invitrogen; 26-well, Bio-Rad Laboratories), and transferred to PVDF (EMD Millipore) at constant 0.5A for 4 h. The PVDF membrane was incubated with antibodies (primary overnight at 4°C h and secondary for 1 h at RT), and bound antibodies were visualized by the enhanced chemiluminescence method (Thermo Fisher Scientific) on a ChemiDoc Imaging System (Bio-Rad Laboratories, Hercules, CA) with ImageLab software v4.0 (Bio-Rad Laboratories).

### Liposome Binding Assay

The liposome binding assay was performed as described for yeast COPII proteins (44, 45) using 10% cholesterol major-minor mix liposomes with Texas Red™ DHPE (Thermo Fisher Scientific) for visualization. Liposomes were extruded through a polycarbonate filter with 100nm pore size (Whatman). Following a 30 min incubation at 37°C, the protein-liposome mixture was diluted into 2.5 M sucrose in HKM buffer to a final concentration of 1 M sucrose. The sample was overlayed with 100μl 0.7 M sucrose and then 20 μl HKM and separated by centrifugation at 391,000 × g for 4 h at 4°C.

### Liposome Aggregation Assay

For visualization by microscopy, each 50μl reaction contained 2μg SAR1 and ∼30k particles/μl liposomes in HKM buffer. Samples were incubated overnight at 37°C and directly pipetted onto a coverslip and imaged with Zeiss Axiovision using the Texas red fluorescent filter. For Nanoparticle tracking analysis, 50μl reaction contained 1μg SAR1 and ∼30k particles/μl liposomes in HKM buffer incubated 3 hours at 37°C. Samples were diluted 1:1000 before analysis.

### GTPase Activity Assay

The tryptophan fluorescence GTPase activity assay was performed at 37°C as described (22–24), using a stirred-cell cuvette. In HKM buffer, we added soluble SAR1B to a final concentration of 1.33 μM and where indicated SEC31 active fragment (24) (2 μM). Five min later, GTP was added to 30 μM. After exchange of GDP for GTP was complete (∼10 min), SEC23-SEC24D complex was added to 250 nM.

### Nanoparticle tracking analysis

Sizes of vesicles budded in vitro were estimated using the NanoSight NS300 instrument equipped with a 405-nm laser (Malvern Instruments, Malvern, United Kingdom). Particles were analyzed in the scatter mode without a filter. Silica 100-nm microspheres (Polysciences, Warrington, PA) were analyzed to check instrument performance and determine the viscosity coefficient of B88. Aliquots (20μl) of vesicles were collected from the top of the flotation gradient as described in the vesicle budding reaction section and diluted 50x with 980 µl filtered B88 (0.02 µm; Whatman). The samples were automatically introduced into the sample chamber at a constant flow rate of 50 (arbitrary manufacturer unit, ∼10 µl/min) during five repeats of 60-s captures at camera level 11 in scatter mode with Nanosight NTA 3.1 software (Malvern Instruments). The particle size was estimated with detection threshold 5 using the Nanosight NTA 3.1 software, after which “experiment summary” and “particle data” were exported. Particle numbers in each size category was calculated from the particle data, in which “true” particles with track length >3 were pooled, binned, and counted with Excel (Microsoft).

### Generation of CRISPR/Cas9 KO cell lines

McArdle-RH7777 cells were transfected with a pX330 vector-derived plasmid(46) containing the targeting sequence from SAR1B (GATGTAGTGTTGGGACGTGCTGG), and a PGK promotor-driven Venus construct (reconstructed by Liangqi Xie from Robert Tjian laboratory at UC Berkeley). After a 24-h transfection, FACS sorting was performed to inoculate single transfected cells in each well of 96-well plates. After 2 weeks, single colonies were expanded and validated by immunoblot and DNA sequencing of the targeted area. Validated positive colonies were employed for the experiments.

### Statistical Analysis

Data in bars represent average ± s.d. Statistical analyses on qualitative data were performed using ANOVA followed by post-hoc Holm p-value adjustment. Statistical analyses on categorical data were performed using Chi-square followed by post hoc pairwise Fisher’s exact test with Holm p-value adjustment using R statistical package.

### Cloning and expression of SAR1 for crosslinking assay

Unnatural amino acids were incorporated into either Sar1A or Sar1B at the noted positions using the following method. pEVOL-pBpf, coding for the tRNA synthetase/tRNA pair, was used for the *in vivo* incorporation of the p-benzoyl-l-phenylalanine into proteins at the position of an in frame amber stop codon (TAG)(47, 48). The mutant Sar1 paralogs, containing TAG codons, were coded for in the pGEX-2T vector. The noted plasmids were cotransformed into DH10B competent cells in a pulser cuvette. The cells were plated and grown overnight at 37°C on LB agar containing chloramphenicol and ampicillin. Colonies were picked and grown at 37°C in LB containing Amp/Cam to an OD of ∼0.4. The temperature was changed to 25°C and protein expression was induced with 1mM IPTG and 0.02% arabinose in the presence of 1mM p-benzoyl-l-phenylalanine. Cells were harvested after 16 h and the protein was purified as previously described (49). The proteins were analyzed by ESI-MS to verify unnatural amino acid incorporation.

### Photoactivated Crosslinking

Photocrosslinking was performed by incubating; Sar1A or B [1µg], 1mM GTP or GTPγS, guanosine 5’-O-[gamma-thio]triphosphate), 2.5 µg of Sec23A/Sec24D and 2.5 µg of Sec13/31A 1.7 µg of liposomes at 32°C for 30 min. Samples were placed in a 96 well flat bottom plate and were irradiated using a handheld ultraviolet lamp (∼360 nm) at 4°C for 5 min. The samples were removed from the wells and resolved on an SDS/PAGE gel and transferred to nitrocellulose and detected by immunoblotting with an antibody recognizing Sec31A.

## Acknowledgments

We thank the staff at the University of California, Berkeley, shared facilities, including Alison Killilea (Cell Culture Facility). We thank Jeremy Thorner for providing the fluorometer and Shawn Shirazi for 3D printing that was essential for the setup. R.S. is supported as an Investigator of the Howard Hughes Medical Institute and the University of California, Berkeley, Miller Institute of Science. D.M. was supported in part by National Institutes of Health grant #11287155.

## Abbreviations and nomenclature

COPII: coat protein complex II
CMRD: chylomicron retention disease
GEF: guanine exchange factor
af: GTPase activating fragment
GTPa: GTP-adjacent
ER: endoplasmic reticulum
ERES: ER exit site

---

**Supplemental Figure 1:**
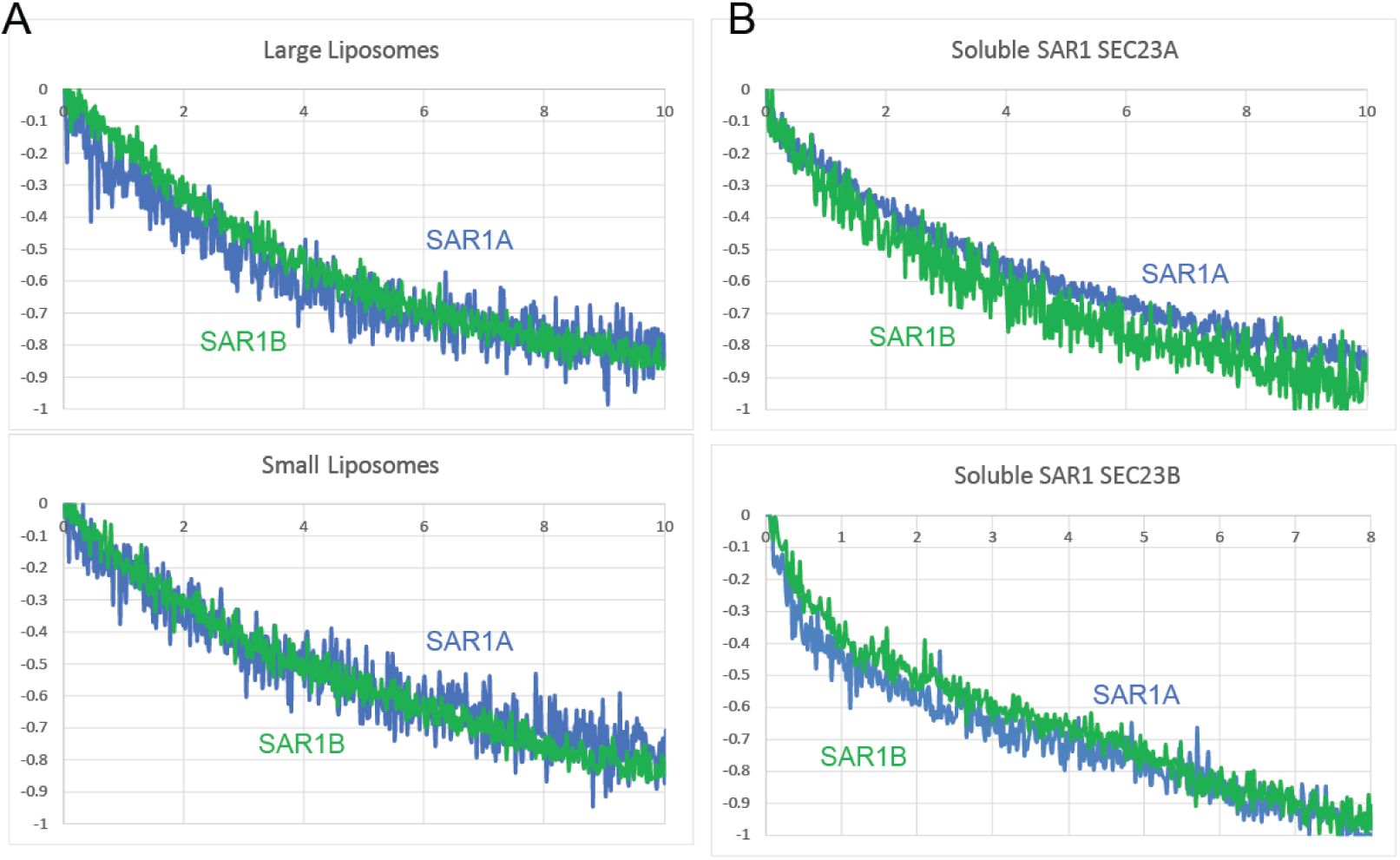
GTPase activity is similar between SAR1A and SAR1B. (A-B) Tryptophan fluorescence assay measuring GTPase activity of SAR1 in presence of SEC31A GTP activating fragment and (A) SEC23A, full length SAR1, and unextruded large liposomes (top) or extruded 100nm liposomes (bottom) or (B) soluble ΔNSAR1 and indicated SEC23 paralog.

